## Supplementary Table for "Domain-PFP: Protein Function Prediction Using Function-Aware Domain Embedding Representations"

**Supplementary Table 1. Training Information of the 12 PLMs in PROBE benchmark.**

| **Model** | **Embedding Size** | **Number of Parameters** | **Training Data** | **Unsupervised tasks used for training** | **Ref.** |
| --- | --- | --- | --- | --- | --- |
| **ProtVec** | 100 | - | Swiss-Prot | word2vec | Asgari & Mofrad, 2015 |
| **LearnedVec** | 128 | - | 524,529 UniProt sequences | doc2vec | Yang et al., 2018 |
| **UniRep** | 5700 | 18.2M | UniRef50 | masked language modelling | Alley et al., 2019 |
| **SeqVec** | 1024 | 93M | UniRef50 | predicting the next amino acid | Heinzinger et al., 2019 |
| **MSA-Transformer** | 768 | 100M | UniRef50 | masked MSA modelling | R. M. Rao et al., 2021 |
| **CPCProt** | 512 | 71M | Pfam | mutual information estimation | Choy et al., 2018 |
| **TAPE-BERT-PFAM** | 768 | 38M | Pfam | masked language modelling | R. Rao et al., 2019 |
| **ProtBERT-BFD** | 1024 | 420M | BFD | masked language modelling | Elnaggar et al., 2021 |
| **ESM-1b** | 1280 | 650M | UniRef50 | masked language modelling | Rives et al., 2021 |
| **ProtXLNet** | 1024 | 409M | UniRef100 | masked language modelling | Elnaggar et al., 2021 |
| **ProtALBERT** | 4096 | 224M | UniRef100 | masked language modelling | Elnaggar et al., 2021 |
| **ProtT5-XL** | 1024 | 3B | UniRef50 | masked language modelling | Elnaggar et al., 2021 |

**Supplementary Table 2. Comparison with 12 PLMs in PROBE benchmark.**

| **Model** | **MF** | **BP** | **CC** | **Avg.** |
| --- | --- | --- | --- | --- |
| **ProtVec** | 0.64 | 0.36 | 0.38 | 0.46 |
| **Learned-Vec** | 0.68 | 0.39 | 0.41 | 0.49 |
| **UniRep** | 0.82 | 0.48 | 0.53 | 0.61 |
| **SeqVec** | 0.89 | 0.60 | 0.61 | 0.70 |
| **MSA-Transformer** | 0.67 | 0.47 | 0.50 | 0.55 |
| **CPCProt** | 0.65 | 0.40 | 0.44 | 0.50 |
| **TAPE-BERT-PFAM** | 0.85 | 0.54 | 0.58 | 0.65 |
| **ProtBERT-BFD** | 0.85 | 0.61 | 0.62 | 0.69 |
| **ESM-1b** | 0.83 | 0.53 | 0.61 | 0.66 |
| **ProtXLNet** | 0.82 | 0.50 | 0.59 | 0.63 |
| **ProtALBERT** | 0.89 | 0.63 | 0.64 | 0.72 |
| **ProtT5-XL** | 0.90 | 0.66 | 0.68 | 0.75 |
| **DomainGO-Prob** | **0.92** | **0.72** | **0.74** | **0.79** |

The value of F1 score is provided for each of the sub-ontologies along with their average value. The scores are collected from Table 2 of Unsal et al., 2022.

**Supplementary Table 3. Overview of NetGO2.0 Benchmark Dataset.**

| **Ontology** | **Number of GO Terms** | **Number of Training Proteins** | **Number of Validation Proteins** | **Number of Test Proteins** |
| --- | --- | --- | --- | --- |
| **MF** | 6,854 | 62,646 | 1,128 | 505 |
| **BP** | 21,814 | 89,828 | 1,124 | 491 |
| **CC** | 2,880 | 81,377 | 1,359 | 268 |

The NetGO2.0 benchmark adopts a timeline-based approach similar to CAFA. Experimentally annotated proteins were obtained from GOA and UniProtKB, and were split into training, validation, and test sets as follows:

1. Training proteins: All proteins annotated on or before December 2018.
2. Validation proteins: Proteins annotated from January 2019 to January 2020.
3. Testing proteins: Proteins annotated between February 2020 and October 2020.

Only GO terms supported by evidence codes EXP, IDA, IPI, IMP, IGI, IEP, TAS, and IC were considered, and the test set comprised proteins from the 17 species of the CAFA4 challenge. In accordance with CAFA guidelines, proteins annotated with only the GO term "protein binding" (GO:0005515) were excluded.

The NetGO2.0 benchmark dataset used in our work used the files that are provided in the google drive url (<https://drive.google.com/drive/folders/1wSS-R335UcNMToMskx3dE4XcTaLvCAOc>) in the NetGO2.0 paper.

But we used the NetGO2.0 dataset that was split into training, validation, and test sets in the DeepGOZero paper by the Hoendorf lab, which they made available at (<https://deepgo.cbrc.kaust.edu.sa/data/deepgozero/data-netgo.tar.gz>). They made this split by following exactly the same procedure described in the NetGO2.0 paper. A practical issue of the NetGO2.0 benchmark dataset was that the data split into training, validation, and test set was only verbally described in their NetGO2.0 paper, but the actual data splits were not provided. Therefore, we used the data splits by the Hoendorf lab.

**Supplementary Table 4. Overview of CAFA3 Benchmark Dataset.**

| **Ontology** | **Number of GO Terms** | **Number of Training Proteins** | **Number of Test Proteins** |
| --- | --- | --- | --- |
| **MF** | 677 | 36,110 | 1,137 |
| **BP** | 3,992 | 53,500 | 2,392 |
| **CC** | 551 | 50,596 | 1,265 |

The CAFA3 challenge organizers developed a training dataset with 66,841 protein sequences and their experimentally annotated functions (EXP, IDA, IPI, IMP, IGI, IEP, TAS, IC evidence codes). The training dataset was released in September 2016, which we used to train our model. A test dataset was provided to evaluate the models, which contains ‘no-knowledge’ proteins that were annotated experimentally between September 2016 and November 2017.

**Supplementary Table 5. Comparison with structure-based function prediction methods**

| **Method** | **F_max_** | | | **AUPR** | | |
| --- | --- | --- | --- | --- | --- | --- |
|  | **MF** | **BP** | **CC** | **MF** | **BP** | **CC** |
| Naive | 0.156 | 0.244 | 0.318 | 0.075 | 0.131 | 0.158 |
| BLAST | 0.498 | 0.400 | 0.398 | 0.120 | 0.120 | 0.163 |
| DeepGO | 0.359 | 0.295 | 0.420 | 0.368 | 0.210 | 0.302 |
| DeepFRI | 0.542 | 0.425 | 0.424 | 0.313 | 0.159 | 0.193 |
| GAT-GO | 0.633 | 0.492 | 0.547 | 0.660 | 0.381 | 0.479 |
| Domain-PFP | 0.662 | 0.519 | 0.532 | 0.653 | 0.503 | 0.510 |
| Domain-PFP + BLAST | **0.746** | **0.611** | **0.605** | **0.754** | **0.606** | **0.612** |

We retrained Domain-PFP on the same dataset used in the DeepFRI (Gligorijević et al., 2021) and GAT-GO (Lai & Xu, 2022) papers. Results of Naïve to DeepFRI are taken from the GAT-GO paper.

The top three scores for each of the metrics are highlighted by bold font, double underline, and single underline, respectively.

**Supplementary Figure 1. Selection of K in the KNN model of Domain-PFP.**

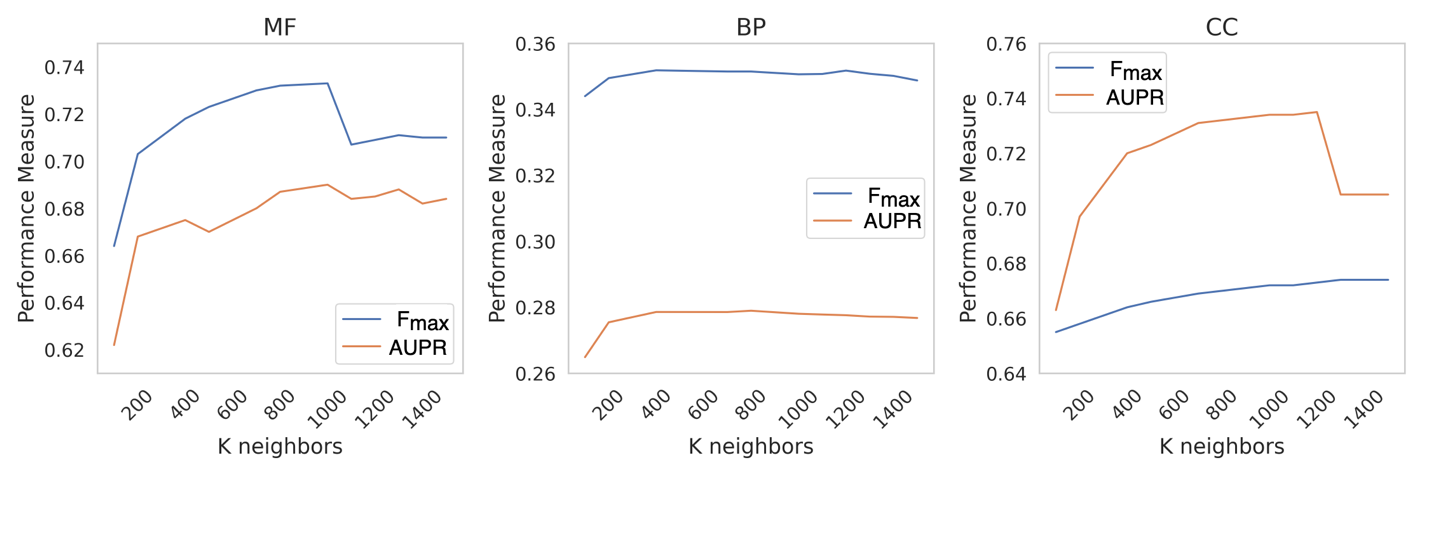

The value of of K for the KNN model was selected by observing the performance for different choices of K on the validation split of the NetGO benchmark dataset. The values of K were selected in the range from 100 to 1500 at an interval of 100 and for each value of K the F_max_ and AUPR scores on the validation data were computed.

We sought to select a value of K that manages to maximize both F_max_ and AUPR scores with more importance put on F_max_. For MF, it was observed that the highest value of F_max_ was obtained with K=1000, after which there was a sharp decline of both F_max_ and AUPR. Thus, we selected K=1000 for MF. For BP, the change in performance for the different values of K was almost minute. We chose K= 800 because the performance was marginally better than the other values for K. For CC, similar to MF, the highest value of F_max_ was observed at K=1200, which dropped sharply with larger K. Although AUPR kept on improving even after K=1200, we choose K=1200 because the increment in AUPR was minor in comparison with the sharp drop in F_max_.
